# Macrophages expressing chimeric cytokine receptors have an inflammatory phenotype and anti-tumoral activity upon IL-10 or TGFβ stimulation

**DOI:** 10.1101/2024.10.01.615826

**Authors:** Sabrina Traxel, Florian Schmidt, Corina Beerli, Dinh-Van Vuong, Roberto F. Speck, Simon Bredl

## Abstract

**Background:** Triple-negative breast cancer (TNBC) is a highly aggressive subtype of breast cancer that lacks hormone receptors and HER2 amplification, making it unresponsive to standard hormone or HER2-targeted therapies. Although immune checkpoint inhibitors (ICIs) have shown promise in tumors with high lymphocyte infiltration, their efficacy remains limited in tumors with minimal lymphocyte infiltration. To overcome this challenge, we developed a novel approach to reprogram tumor-associated macrophages (TAMs) from an immunosuppressive to an inflammatory phenotype within the tumor microenvironment (TME), aiming to enhance therapeutic outcomes in TNBC.

**Methods:** We designed a chimeric cytokine receptor (ChCR) that triggers STAT1 signaling upon stimulation with IL-10 or TGFβ, cytokines prevalent in the TME that typically drive immunosuppression. Human primary macrophages were transduced with a lentiviral vector expressing the ChCR. Changes in their phenotype, secretome, and transcriptome were analyzed following stimulation. The anti-tumoral activity of these reprogrammed macrophages was assessed using co-culture assays with 3D TNBC spheroids.

**Results:** ChCR-expressing macrophages showed robust STAT1 activation in response to IL-10 or TGFβ stimulation, resulting in an inflammatory phenotype similar to IFNγ activation, as confirmed by phenotypic markers, and transcriptomic profiling. These ChCR-stimulated macrophages demonstrated significant anti-tumoral effects in 3D TNBC spheroids. Moreover, ChCR stimulation led to the upregulation of *CXCL9* and *CXCL10*, chemokines essential for lymphocyte recruitment, and genes associated with good response to ICIs.

**Conclusion:** We successfully engineered ChCRs that reprogram TAMs within an IL-10- and TGFβ-rich environment, inducing an inflammatory phenotype and anti-tumoral activity. The expression of *CXCL9* and *CXCL10* further supports lymphocyte recruitment, potentially facilitating greater lymphocyte infiltration and disrupting the immunosuppressive TME. This approach offers a promising strategy to improve immunotherapy outcomes in TNBC patients, particularly those with low immune cell infiltration, thereby addressing a critical unmet clinical need.

**What is already known on this topic**

Low T cell infiltration in triple negative breast cancer is associated with bad prognosis and non-responsiveness to immune checkpoint inhibitor in combination with chemotherapy. The presence of macrophages with an inflammatory phenotype is associated with T cell infiltration and better prognosis. The tumor microenvironment (TME), however, is rich in immunosuppressive cytokines like IL-10 and TGFβ, rendering macrophages immunosuppressive. Thus, reprogramming macrophages in the TME could benefit patients with low T cell infiltration.

**What this study adds**

We have designed a chimeric cytokine receptors (ChCR) that bind IL-10 or TGFβ and induces an IFNγ-like inflammatory signaling and thus utilizes the local presence of immunosuppressive cytokines to reprogram macrophages. ChCR expressing macrophages show an inflammatory phenotype in the presence of immunosuppressive cytokines IL-10 or TGFβ and have an anti-tumoral activity *in vitro*.

**How this study might affect research, practice or policy –**

Our study is a starting point to explore ChCR-expressing macrophages as an adoptive cell therapy. Such therapy has the potential to attract T cells into the tumor and thereby boost the adaptive anti-tumoral immune response and response to immune checkpoint inhibitors. Moreover, our data suggest that such genetically-engineered macrophages have direct anti-tumoral activity.

## INTRODUCTION

Triple negative breast cancer (TNBC) is an aggressive subtype of breast cancer that lacks both hormone receptor expression and HER2 amplification and is therefore not amenable to hormone or HER2 targeted therapies (reviewed^1^). Current first line treatment for both high-risk early-stage TNBC and advanced stage, PD-L1 positive (combined positive score ≥10) TNBC is the immune checkpoint inhibitor (ICI) pembrolizumab in combination with chemotherapy. Early stage TNBC has a good prognosis with 86.6% overall survival at 60 months when treated neoadjuvant with pembrolizumab and chemotherapy.^2^ In patients with advanced stage TNBC the pembrolizumab and chemotherapy combination minimally improves the median overall survival to 23 months, compared to 16.1 months in the placebo and chemotherapy group.^3^ Thus, there is clearly an unmet need for patients with advanced TNBC.

Adoptive cell therapy using chimeric antigen receptor (CAR) T cells has achieved remarkable success in hematological malignancies^4^ and there was great hope that CAR T cell therapy would also change the therapeutic landscape for solid tumors. To date, CAR T cells had only limited success in solid tumors.^5^ Several phase I clinical trials of CAR-based immunotherapies for TNBC are currently underway.^6^ While we eagerly await the first results of adoptive CAR T cell therapies in TNBC, alternative innovative treatments must be sought.

A critical determinant of disease progression of solid tumors including TNBCs is their immunophenotype, which is based on the number and localization of CD8+ T cells in the tumor microenvironment (TME).^7–9^ Three main immunophenotypes have been defined: hot (high number of intratumoral T cells), excluded (T cells restricted to the tumor margin) or cold (lack of T cells) (reviewed^10^). Patients with a hot TME, including patients suffering from TNBC, have a better prognosis and show better response to ICI treatment than patients with an excluded or cold TME. ^7^ ^9^ ^11^ ^12^ In TNBC, 42-54% of patients have a tumor with excluded or cold immunophenotype.^7^ ^9^ ^13^ Therefore, shifting a cold into a hot TME is of great therapeutic interest.

Monocytes are actively recruited into the TME and there, differentiate to tumor associated macrophages (TAMs), which, together with other myeloid cells, they constitute up to 30% of the tumor mass.^14^ ^15^ Macrophages can adopt a wide range of phenotypes depending on the stimuli received and on the microenvironment.^16^ Consequently, their activities include immunosuppression, tissue repair, phagocytosis, inflammation, and cytotoxicity. *In vitro*, we can model extreme macrophage phenotypes: Inflammatory macrophages are induced by pro-inflammatory signals like IFNγ and/or LPS, whereas immunosuppressive macrophages are induced by immunosuppressive cytokines like IL-4, IL-13, IL-10, or TGFβ. Notably, these extreme phenotypes cannot describe the complexity of phenotypes in macrophages found *in vivo*, where features of both extremes can be found within the same cell (reviewed^17^). The presence of IL-10 and TGFβ in the TME of solid tumors, such as in TNBC^18–20^, however, pushes TAMs towards an immunosuppressive phenotype. These pro-tumoral TAMs inhibit the immune system, support tumor growth, angiogenesis, and tumor cell migration (reviewed^21^). In contrast, inflammatory macrophages have anti-tumoral activity by attracting T cells, by activating adaptive immune response, and by their direct tumoricidal effects.^22^ ^23^ (reviewed^24^) In fact, tumors with high number of inflammatory macrophages are more likely to have a hot TME, a better prognosis and better response to ICIs.^7^ ^22^ ^25^ ^26^ Thus, reprogramming TAMs towards an inflammatory phenotype is a hot topic in the field of cancer immunotherapy (reviewed^17^).

We believe that macrophages should be reprogrammed to switch to inflammatory and anti-tumoral macrophages locally in the TME. Systemically activated macrophages may be retained in various tissues, unable to reach their destination. Furthermore, local activation would be less likely to have off-target effects. Therefore, we developed chimeric cytokine receptor (ChCR) expressing macrophages that utilize immunosuppressive cytokines, prevalent in the TME, to switch to inflammatory macrophages. Specifically, we engineered macrophages to express a ChCR that binds either IL-10 or TGFβ and induces an IFNγ signaling.

## METHODS

### Ethics statement

Monocytes used were obtained from buffy coats, anonymously provided by the Blood Donation Service Zurich, Swiss Red Cross, Schlieren. Written consent for the use of buffy coats for research purposes was obtained from blood donors by the Blood Donation Centre.

### Cell lines

HEK293T (ATCC) were maintained in DMEM (high glucose, Sigma-Aldrich, D6429) with 10% heat-inactivated FBS (Gibco, 10270-106) (hereafter called D10). HEK-Blue^TM^ IFN-γ cells (Invivogen, hkb-infg)) were maintained in D10 with 100µg/ml Zeocin (Invivogen, ant-zn-1) and 30µg/ml Blasticidin (Invivogen, ant-bl-1). MDA-MB-231 (ATCC, HTB-26) were maintained in RPMI-1640 (Sigma-Aldrich, R8758) with 10% heat-inactivated FBS (hereafter called R10). BT-549 (ATCC, HTB-122) were maintained in R10 with 10ug/ml insulin (Sigma-Aldrich, I9278). THP-1 (ATCC) were maintained in R10 with 2-mercaptoethanol (Gibco, 31350010). MRC-5 (ATCC, CCL-171) were maintained in EMEM (ATCC, 30-2003) with 10% heat-inactivated FBS. All cells were maintained at 5% CO_2_ and 37°C in a humidified incubator.

### Monocyte derived macrophages

Peripheral blood mononuclear cells were isolated from buffy coats of healthy donors (Blood Donation Service Zurich, Swiss Red Cross, Schlieren) using density gradient centrifugation on lymphoprep (Axon Lab, 12053230). Monocytes were isolated using CD14 microbeads (Miltenyi Biotec, 130-050-201) and plated at 150’000 cells/cm^2^ in F+ high attachment plates (Sarstedt) in RPMI-1640 with Penicillin-Streptomycin (P/S, Gibco, 15140-122). Two hours after plating, medium was replaced with RPMI-1640 with P/S and 10% human AB serum (Sigma Aldrich, H6914), hereafter called RH10. Monocytes were differentiated for 6-11 days with medium change every 2-3 days. In some experiments, macrophages were stimulated for additional 48h with IFNγ (Biolegend, 570204), IL-10 (Biolegend, 571004) or TGFβ1 (Peprotech, 100-21) at indicated concentrations.

### Plasmid construction

To generate ChCR plasmids, amino acid sequences of the human IL10RA, IL10RB, TGFβR1, TGFβR2, IFNGR1 and IFNGR2 receptor were obtained from UniProt (Supplementary Table 1). The ChCR were constructed as follows: The extracellular domains of IL10RA, IL10RB, TGFβR1, or TGFβR2 were fused to the transmembrane and extracellular domain of the IFNGR1 or IFNGR2, including a glycine-serine linker for flexibility.^27^ As an N-terminal signal peptide, we used the one from the corresponding intracellular IFNGR. All variants were analyzed with DeepTMHMM to verify *in silico* signal peptide cleavage, transmembrane domains and surface expression.^28^ Sequences were ordered codon-optimized at GenScript Biotech Corporation. For expression of single chains, ChCR sequences were sub-cloned into a pCWX Dest lentiviral vector (Gift from Patrick Salmon) under the control of an elongation factor 1 alpha core promoter (EF1α). To simultaneously express both chains, ChCR sequences were subcloned into a pCWX Dest lentiviral vector separated by a P2A sequence and under the control of a spleen focusLforming virus (SFFV) promoter. All lentiviral vectors also encode for a truncated nerve growth factor receptor (tNGFR) driven by a human phosphoglycerate kinase (hPGK) promoter as a reporter for transduction. Mock lentiviral vector only encodes for tNGFR.

To generate the Luciferase-EGFP plasmid, luciferase and EGFP sequences, separated by a T2A sequence, were ordered as gene blocks and sub-cloned into a pCWX Dest lentiviral vector under the control of EF1α promoter.

### Lentivirus production and titration

Lentiviral vectors were produced in HEK293T. Briefly, 13 million cells were plated in T150 flasks. The day after, cells were co-transfected using PEI (Polysciences, 23966) with pMD2.G, p8.9NDSB-DPAVDLL^29^ (gift from Jeremy Luban), pcDNA3.1 Vpx239^29^ (gift from Jeremy Luban) and pCWX Dest transfer vector at a molar ratio of 251:107.5:756.1:45. 4h after transfection, medium was changed and supplemented with 5mM sodium butyrate (Sigma Aldrich, B5887). Virus was harvested 52h after transfection and clarified by centrifugation. Lentiviral supernatants were concentrated by ultrafiltration using 100kDa cut-off Amicon filters (Merck Milipore, UFC910024). Titration of lentiviruses was done by transducing HEK293T. Three days after transduction, transduced cells were stained for tNGFR and analyzed by flow cytometry on a LSR II Fortessa (BD Biosciences).

### Stable cell line generation

THP-1 were transduced with lentiviruses at a multiplicity of infection (MOI) 1 by spinoculation (800g, 45min, 32°C) in presence of protamine sulfate (100ug/ml, Sigma-Aldrich, P4020). Transduced cells (tNGFR+) were sorted using MACSelect LNGFR microbeads (Miltenyi Biotec, 130-091-330).

MDA-MB-231 cell line was transduced with the lentiviral construct to express Luciferase and EGFP. Cells were sorted for high EGFP expression on a BD FACSAria III (BD Biosciences).

### Monocyte transduction

To transduce monocytes, lentivirus was added 2h after plating of monocytes in RPMI with P/S and 5% human AB serum at a MOI of 10 or 15 in presence of protamine sulfate (100ug/ml, Sigma-Aldrich, P4020). After overnight incubation, medium was changed to RH10. Transduction efficiency was assessed between day 6 and day 12 of differentiation and was defined as %tNGFR+ cells. In general, macrophages with transduction rates < 65% were excluded from experiments. In early experiments assessing IL-10 ChCR macrophage polarization, transduction rates ≥ 55% were accepted.

### IFN**γ** signaling reporter cells

HEK-Blue^TM^ IFN-γ were transduced with lentiviruses by spinoculation in the presence of 8ug/ml polybrene (SantaCruz, sc-134220). To measure secreted embryonic alkaline phosphatase (SEAP), transduced cells were detached mechanically and seeded in 96-well plate. Cells were incubated with cytokines as indicated for 24h. Thereafter, 30ul of the cell supernatant was incubated with 170ul Quanti-Blue solution (Invivogen, rep-qbs) and incubated for 1h at 37°C. OD was measured at 620nm using Spectramax iD3 plate reader (Molecular Devices).

### Flow cytometry analysis

To assess transduction rate and polarization marker expression, macrophages were harvested for flow cytometry by incubation with 5mM EDTA in PBS on ice for 45min, followed by gentle pipetting. Detached macrophages were first treated with True-Stain FcX (Biolegend, 422301) and thereafter stained with antibodies (Table 1). Stained samples were acquired on LSR II Fortessa (BD Life Sciences). Untransduced or mock transduced cells were used as gating control for tNGFR and IL-10Rα, IL-10Rβ, and TGFβR2, respectively. Data was analyzed using FlowJo v10.10 software (BD Life Sciences). Corresponding isotypes were used to normalize median fluorescence intensities (MFI) by subtraction or as a gating control.

**Table 1:**
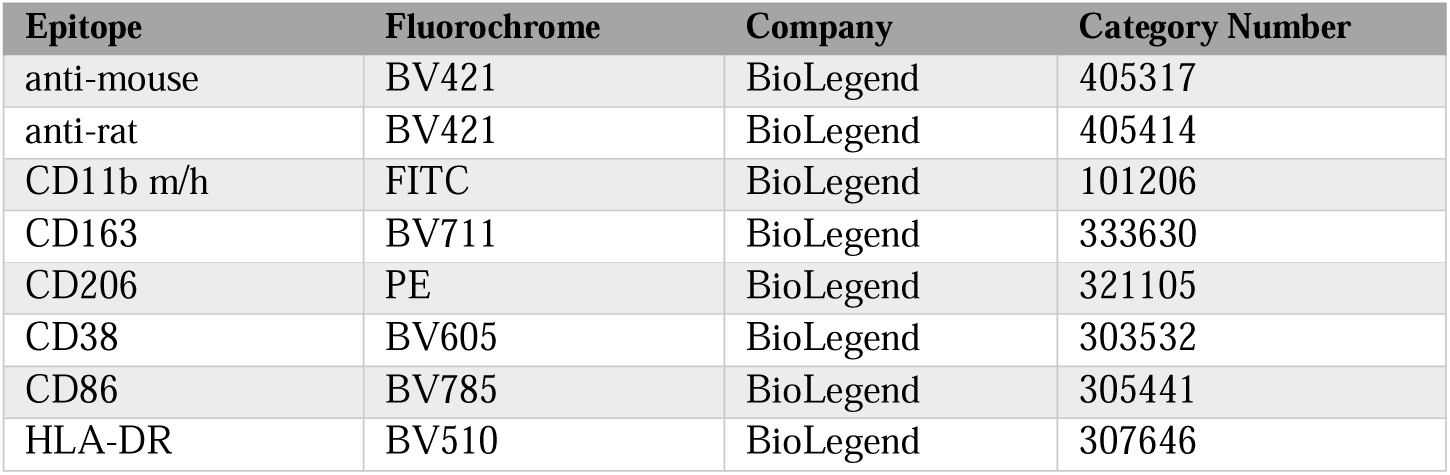
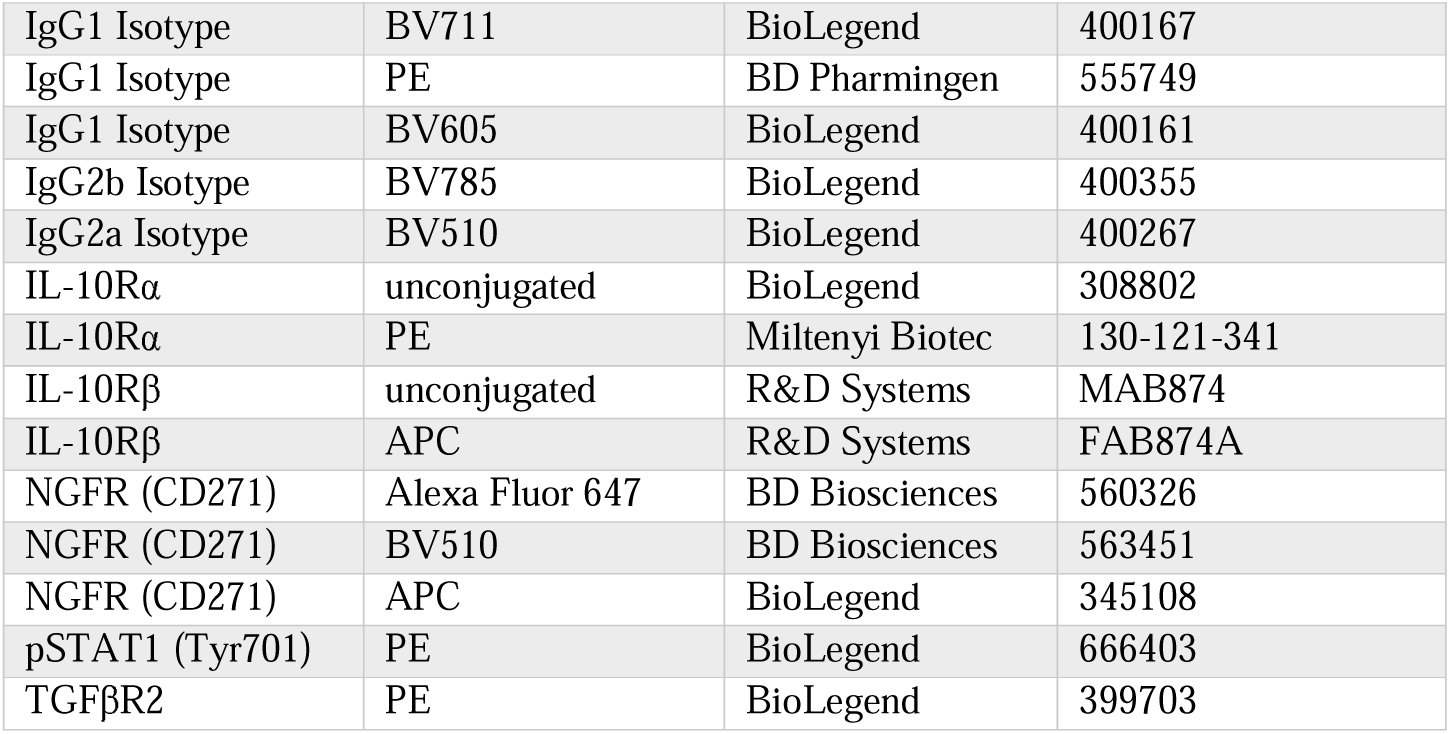
Flow cytometry antibodies.

### STAT1 phosphoflow

THP-1 or human primary macrophages were used for STAT1 phosphoflow experiments. THP-1 were plated in round bottom 96-well plates in R10. Macrophages were detached as described for flow cytometry analysis and plated in round bottom ultra low attachment 96-well plates (Corning, 7007) in RH10. Cells were rested for 1h at 37°C and thereafter stimulated for 30min as indicated. Cells were fixed by adding one volume of 4% PFA (Santa Cruz, sc-281692) and 15min incubation at 37°C. Next, cell surface markers were stained before cells were permeabilized with True-Phos Perm Buffer (Biolegend, 425401). Cells were stained for intracellular pSTAT1 (Tyr701) and analyzed on LSR II Fortessa (BD Biosciences).

### *IL10* and *TGFB1* expression in TNBC

Gene expression data from 195 TNBC patients were obtained from The Cancer Genome Atlas (TCGA). TNBC patients in the TCGA dataset were previously identified by Lehmann et al.^30^ Raw counts were retrieved, preprocessed and normalized using the R package TCGABiolinks (2.33.0) without filtration.^31^ Percentile expression of *IL10* and *TGFB1* across all genes was calculated by dividing the rank expression with the total analyzed genes.

### RNA sequencing (RNAseq)

#### RNAseq and preprocessing

For bulk RNAseq, monocytes were transduced and differentiated into macrophages. Macrophages were stimulated for 48h as indicated. Total RNA was extracted using Quiashredder (Qiagen, 79654)) and RNeasy Mini Kit (Qiagen, 74104) with on column DNA digestion (Qiagen, 79254). RNAseq was performed at the Functional Genomics Center Zurich (FGCZ). The NovaSeq X Plus (Illumina, Inc, California, USA) was used for cluster generation and sequencing according to standard protocol. Sequencing was paired end 150 bp. Between 28 and 86 million reads were generated per sample. RNAseq analysis was performed using the SUSHI framework,^32^ which encompassed the following steps: Read quality was inspected using FastQC,^33^ and sequencing adaptors removed using fastp,^34^ alignment of the RNA-Seq reads using the STAR aligner,^35^ and with the GENCODE human genome build GRCh38.p13 (Release 42) as the reference^36^, the counting of gene-level expression values using the featureCounts function of the R package Rsubread.^37^ Unless stated otherwise, all R functions were executed on R version 4.3.2 (R Core Team, 2020) and Bioconductor version 3.17.

#### RNAseq analysis

Differential expression analysis was performed using the generalized linear model as implemented by the DESeq2 Bioconductor R package.^38^ Principle component analysis (PCA) was performed using the prcomp function from stats R package. For gene set enrichment analysis (GSEA), differentially expressed genes were ranked by log2 fold change. Gene Ontology (GO) GSEA tests were performed using the gseGO function respectively of the clusterProfiler Bioconductor R package.^39^ Hallmark and custom pathways GSEA was performed using the GSEA function of the Bioconductor R package clusterProfiler (4.13.0)^39^ and the hallmark gene sets (MSigDb 2023.2)^40^ on R version (4.4.0).

#### Analysis of previously published scRNAseq data

We compared the transcriptome of ChCR macrophages with previously described scRNAseq data of macrophages from TNBC patients with or without response to anti-PD-L1 atezolizumab.^25^ Using the R package VennDiagram (1.7.3), we determined the overlap of significantly upregulated genes in ChCR macrophages (log2FC>0.5 and false discovery rate (FDR)<0.05) and genes upregulated in baseline tumor macrophages of anti-PD-L1 responders previously published. Moreover, we performed GSEA with costum pathways generated from genes upregulated in baseline tumor macrophages from anti-PD-L1 responders and non-responders, respectively. GSEA was performed using the GSEA function of the Bioconductor clusterProfiler (4.13.0)^39^ on R version (4.4.0).^25^

### CXCL10 ELISA

To analyze CXCL10 secretion, differentiated macrophages were stimulated for 48h as indicated. Supernatants were collected, cell debris removed by centrifugation, and stored at −80°C. CXCL10 in supernatant was measured using the ELISA MAX™ Deluxe Set Human CXCL10 (IP-10) kit (Biolegend, 439904). OD450nm and OD570 as background were measured using Multiskan SkyHigh plate reader (Thermo Scientific^TM^).

### 3D spheroid assay

Heterotypic spheroids were generated by mixing 1000 MDA-MB-231, expressing EGFP and luciferase, and 1500 MRC-5 cells in Akura™ 96 Spheroid Microplates (InSphero, CS-PB15) in R10. Plates were centrifuged at 100g for 5 min to remove air bubbles and incubated at 37°C, 5% CO_2_ overnight. To support the aggregation and maturation of the spheroids, the plates were placed on a tilting stand at a 30° angle. The following day, macrophages were detached as described for flow cytometry analysis, added at a 10:1 ratio (macrophages:MDA-MB-231 at seeding), and stimulated as indicated. Medium and stimuli were replaced every 2-3 days. Fluorescent (EGFP) images were captured using the Cytation 3 (BioTek) with a 4x objective. Images were processed and analyzed with the Gen5 (BioTek) software. The mean fluorescence intensity of the full image was determined and the background of a medium control well subtracted.

### Luciferase assay

As an endpoint quantification of tumor cells in the spheroids, luciferase activity of the remaining MDA-MB-231 was determined at day 6. Spheroids were washed twice with PBS to remove luciferase in the supernatant and left with 50 ul residual PBS per well. 50 ul of BrightGloTM substrate (Promega, E2610) was added per well to lyse remaining cells. The lysate was transferred into a 96-well white plate (Pierce™, 15042) and incubated for minimum 2min. Luciferase activity was determined using GloMax® Explorer luminometer (Promega, GM3500) and background signal from a medium control well was subtracted.

### WST1 assay

To assess the ability of genetically-engineered macrophages to affect TNBC proliferation, macrophages were stimulated for 2d with IFNγ, IL-10 or TGFβ at indicated concentrations. Supernatants were harvested, clarified by centrifugation and stored at −80°C. For WST1 assay, 5’000 BT-549 or 10’000 MDA-MB-231 cells were plated in 96-well plates. The next day, medium was replaced with supernatants from stimulated macrophages or RH10 as untreated control. 72h after addition of supernatants, viability was measured by incubating cells for 1h with cell proliferation reagent WST-1 (Roche, 5015944001). OD440nm and OD650nm as background were measured using Multiskan SkyHigh plate reader (Thermo Scientific^TM^).

### Statistical analysis

The statistical analysis of TCGA and previously published single cell RNAseq data was performed in R (4.4.0). The statistical analysis of bulk RNAseq data was performed using the Sushi framework of the FGCZ.^32^ The remaining statistical analysis was performed using Graphpad Prism 10.2.0 using statistical tests specified in the figure legends.

### Data availability

RNA-Seq data are deposited on the Gene Expression Omnibus database under accession number GSE280810.

## RESULTS

### *IL10* and *TGFB1* are expressed in TNBC

IL-10 and TGFβ1 have previously been shown to be expressed in TNBC patients.^18–20^ To confirm this data, we analyzed publicly available RNAseq data from 195 TNBC patients for *IL10* and *TGFB1* expression. We found that *IL10* and *TGFB1* were expressed among the top 73% (median) and top 90% (median) genes, respectively (figure 1A). Thus, TCGA gene expression analysis confirms previously published results of IL-10 and TGFβ expression in TNBC patients.

**Figure 1.**
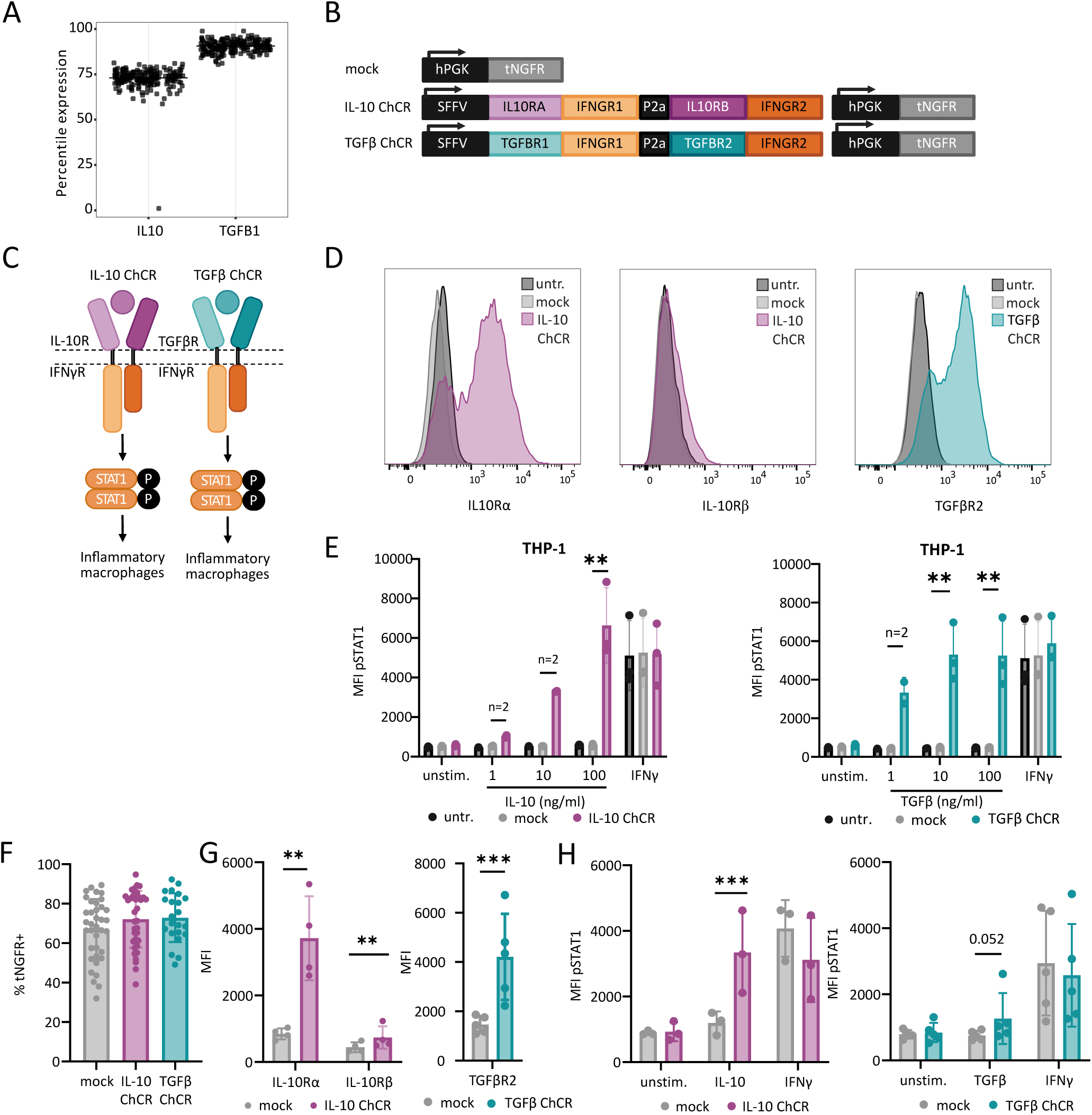
IL-10 / TGFβ stimulation induces STAT1 signaling in chimeric cytokine receptors (ChCR) expressing THP-1 and primary macrophages. A) Expression of *IL10* and *TGFB1* in TNBC. Shown is the percentile expression of *IL10* and *TGFB1* expression across all transcripts in TNBC samples from TCGA database. Each dot represents one patient. n=195 TNBC patients. B) Scheme of lentiviral constructs that were used. C) Scheme of ChCR. Binding of IL-10 or TGFβ to the respective ChCR induces pSTAT1 signaling. C/D) THP-1 were either left untransduced or transduced with mock, IL-10 ChCR or TGFβ ChCR construct. C) Expression of IL-10Rα, IL-10Rβ and TGFβR2 was assessed by flow cytometry. Shown are representative histograms. Cells are gated on tNGFR+ cells. D) THP-1 were either left untreated, stimulated with 1, 10 or 100ng/ml IL-10 or TGFβ, or 5ng/ml IFNγ for 30min. pSTAT1 signaling was assessed by Phosphoflow. Shown are median fluorescence intensities (MFI’s, mean ± SD) in single cells for untransduced or in tNGFR+ cells for mock, IL-10 ChCR or TGFβ ChCR. N=2-3 E/F) Primary human monocytes were either transduced with mock, IL-10 ChCR or TGFβ ChCR construct. E) Transduction efficiency (tNGFR+) was assessed by flow cytometry upon differentiation to macrophages. Shown are % tNGFR+ cells (mean ± SD). N=14-33 F). Expression of IL-10Rα, IL-10Rβ and TGFβR2 was assessed by flow cytometry. Shown are MFIs gated on tNGFR+ cells (mean ± SD) from 4-5 healthy donors from 3-4 independent experiments G) Transduced macrophages were either left untreated or stimulated as indicated for 30min. pSTAT1 signaling was assessed by Phosphoflow. Shown are pSTAT1 MFIs in tNGFR+ cells (mean ± SD) from 4 donors in 4 independent experiments (IL-10 ChCR) or 5 donors in 3 independent experiments (TGFβ ChCR). p-values were calculated by ratio paired t-tests. * p ≤ 0.05, ** p≤0.01, *** p≤0.001, **** p≤0.00001

### IL-10 and TGF**β** ChCR induce STAT1 signaling

To convert IL-10 or TGFβ signals into IFNγ signals, we designed two ChCR, which consist of the extracellular domains of IL-10 receptor or TGFβ receptor, respectively, fused to the intracellular domains of the IFNγ receptor (figure1B and C). To control for transduction efficacy, we also incorporated the tNGFR in the lentiviral vector.

In a first instance, we verified IFNγ signaling through the IL-10 ChCR in HEK-Blue IFNγ reporter cell line. In the HEK-Blue IFNγ reporter cell line, IFNγ-induced STAT1 signaling leads to expression of SEAP, which can be measured in the supernatant using a colorimetric assay. We stably expressed either the full IL-10 ChCR or the individual components, i.e. IL-10RA fused to IFNGR1 and IL-10RB fused to IFNGR2 in the HEK-Blue IFNγ reporter cells (online supplemental figure S1A). All cell lines were transduced at >90% efficiency (data not shown) and ChCR expression was confirmed by flow cytometry (online supplemental figure S1B). We found that IL-10 induced IFNγ signaling in a dose-dependent manner when the full IL-10 ChCR was expressed (online supplemental figure S1C). Surprisingly, IL-10RA-IFNGR1 alone triggered moderate IFNγ signaling, but both chains of the ChCR were necessary to achieve maximum STAT1 signaling. We also verified that the expression of the full TGFβ ChCR in HEK-Blue IFNγ reporter cell line induced STAT1 signaling (data not shown). Next, we proceeded to the characterization of both ChCRs in the monocyte cell line THP-1, which more closely approximates the properties of primary human macrophages. We found a prominent upregulation of IL-10Rα and TGFβR2 and a moderate one of IL10Rβ using flow cytometry (figure1D). We tested several TGFβR1 antibodies, all suffered from non-specificity and therefore we do not have expression data for TGFβR1. Since TGFβR1 is on the first position of the polycistronic vector, and TGFβR2 expression is detectable, we assume that TGFβR1 is expressed as well. Importantly, IL-10 and TGFβ induced a dose-dependent activation of STAT1 in IL-10 or TGFβ ChCR-expressing THP-1 cells, respectively (figure 1E). As expected, untransduced and mock transduced THP-1 cells only showed pSTAT1 signaling upon IFNγ stimulation. To investigate the ChCRs in primary macrophages, we first optimized the protocol for transducing monocytes. We achieved a transduction efficiency of 70.2 ± 14.5% (mean±SD) across all vectors (figure 1F). Similar to ChCR expression in THP-1, we saw a very prominent upregulation of IL-10Rα and TGFβR2 and a lesser one of IL-10Rβ (figure 1G). IL-10 induced STAT1 activation to a similar level like IFNγ, whereas TGFβ induced only a little, but consistent STAT1 activation in the macrophages expressing the ChCRs (figure1H). Thus, both IL-10 and TGFβ ChCR variants trigger the IFNγ signaling pathway.

### ChCR expression induces an inflammatory phenotype upon IL-10 / TGF**β** stimulation

We next assessed whether triggering the ChCRs induces an inflammatory phenotype in primary human macrophages. We focused on the expression of polarization markers typically upregulated on macrophages either upon IFNγ (i.e., HLA-DR, CD38, CD86) or IL-10 stimulation (i.e., CD163). Importantly, both, IL-10 and TGFβ induced the expression of HLA-DR, CD38 and CD86 in the macrophages expressing the corresponding ChCRs (figure 2A and B). Moreover, IL-10 blocked the upregulation of CD163 in IL-10 ChCR transduced macrophages, while IL-10 massively increased its expression in the controls (figure 2A). Notably, CD163 is upregulated in response to IL-10 but not to TGFβ and therefore we did not measure CD163 expression in TGFβ ChCR-expressing macrophages. Overall, we found that, ChCR-expressing macrophages indeed show a phenotype similar to IFNγ stimulated macrophages upon stimulation with IL-10 or TGFβ.

**Figure 2.**
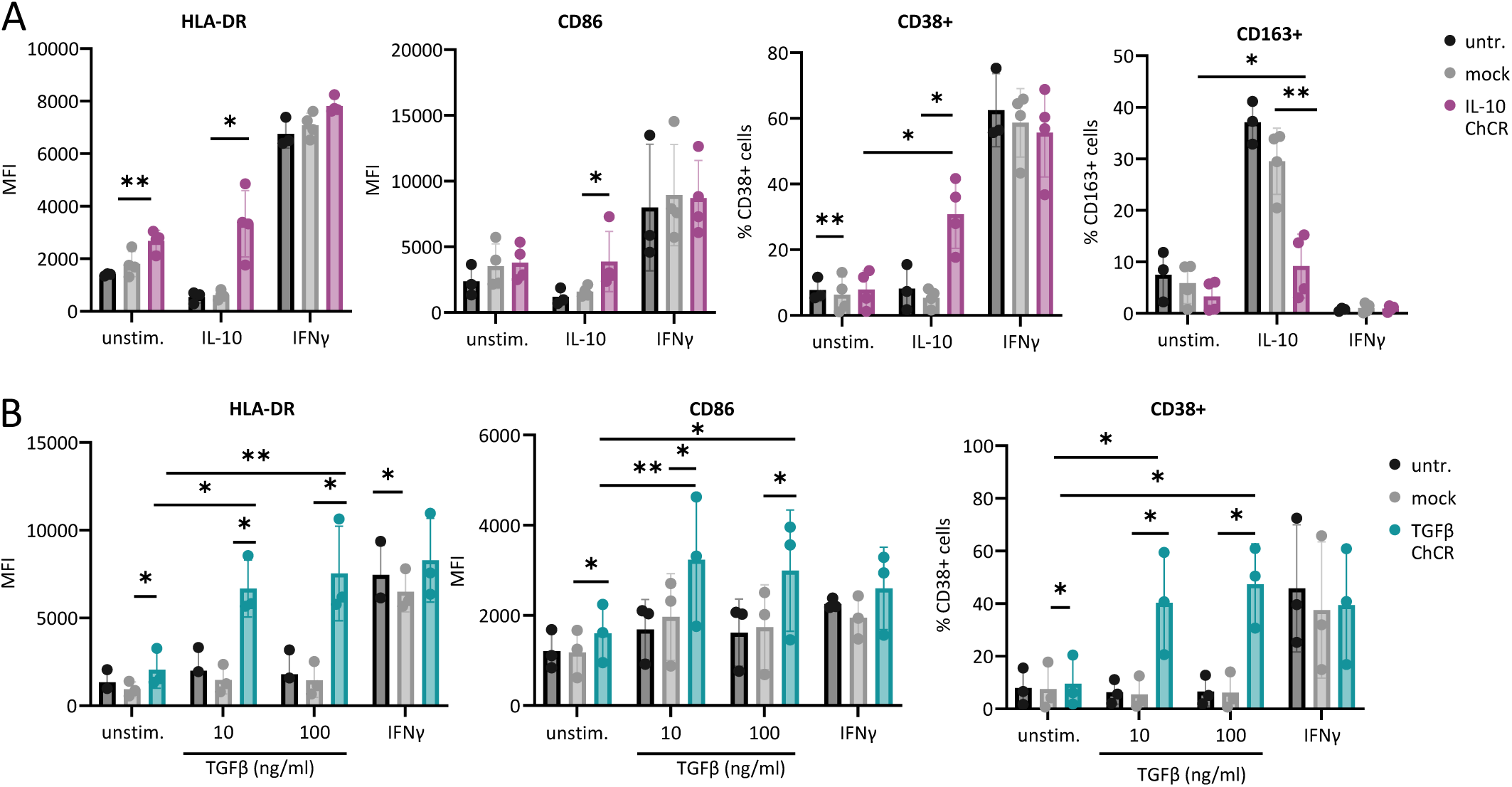
ChCR-expressing macrophages have an inflammatory phenotype upon IL-10 or TGFβ stimulation. A/B) Primary human monocytes were either left untransduced or transduced with mock, IL-10 ChCR or TGFβ ChCR construct. After differentiation to macrophages, cells were either left unstimulated or stimulated with 100ng/ml IL-10, 10 or 100ng/ml TGFβ, or 10ng/ml IFNγ. Expression of surface marker was assessed by flow cytometry. A) Shown are median fluorescence intensities (MFIs, mean ± SD) after subtracting isotype MFI or % CD38+ or % CD163+ gated on CD11b+ cells from 3-4 healthy donors in 4 independent experiments. B) Shown are MFI (mean ± SD) after subtracting isotype MFI or % CD38+ gated on CD11b+ (untransduced) or NGFR+ (transduced) cells from 3 healthy donors in 2 independent experiments. For MFIs, p-values were calculated using ratio paired t-tests. For % positive cells, p-values were calculated using paired t-tests. * p ≤ 0.05, ** p≤0.01, *** p≤0.001, **** p≤0.00001

### ChCR-expressing macrophages have an IFN**γ**-like transcriptome

Next, we performed RNAseq to extend the characterization of ChCR-expressing macrophages. PCA showed that ChCR-expressing macrophages stimulated with their corresponding ligand clustered together with IFNγ stimulated-macrophages, indicating an IFNγ-like transcriptome profile induced by stimulation of the ChCRs (figure 4A). As expected, differential gene expression analysis of ChCR-expressing macrophages showed the upregulation of many IFNγ stimulated genes upon IL-10 or TGFβ stimulation when compared to mock transduced macrophages stimulated equally (figure 3B). Similarly, gene set enrichment analysis (GSEA) of hallmark gene sets showed that IFNγ stimulated genes, among other *CXCL9, CXCL10, and CIITA,* were significantly enriched in ChCR-expressing macrophages upon stimulation (figure 3C). Moreover, GSEA of gene ontology biological processes (GO:BP) showed enrichment of genes associated with antigen presentation (MHC-II HLA genes) and several immune related pathways (figure 3D). Thus, we could confirm that ChCR-expressing macrophages have an IFNγ-like transcriptome.

**Figure 3.**
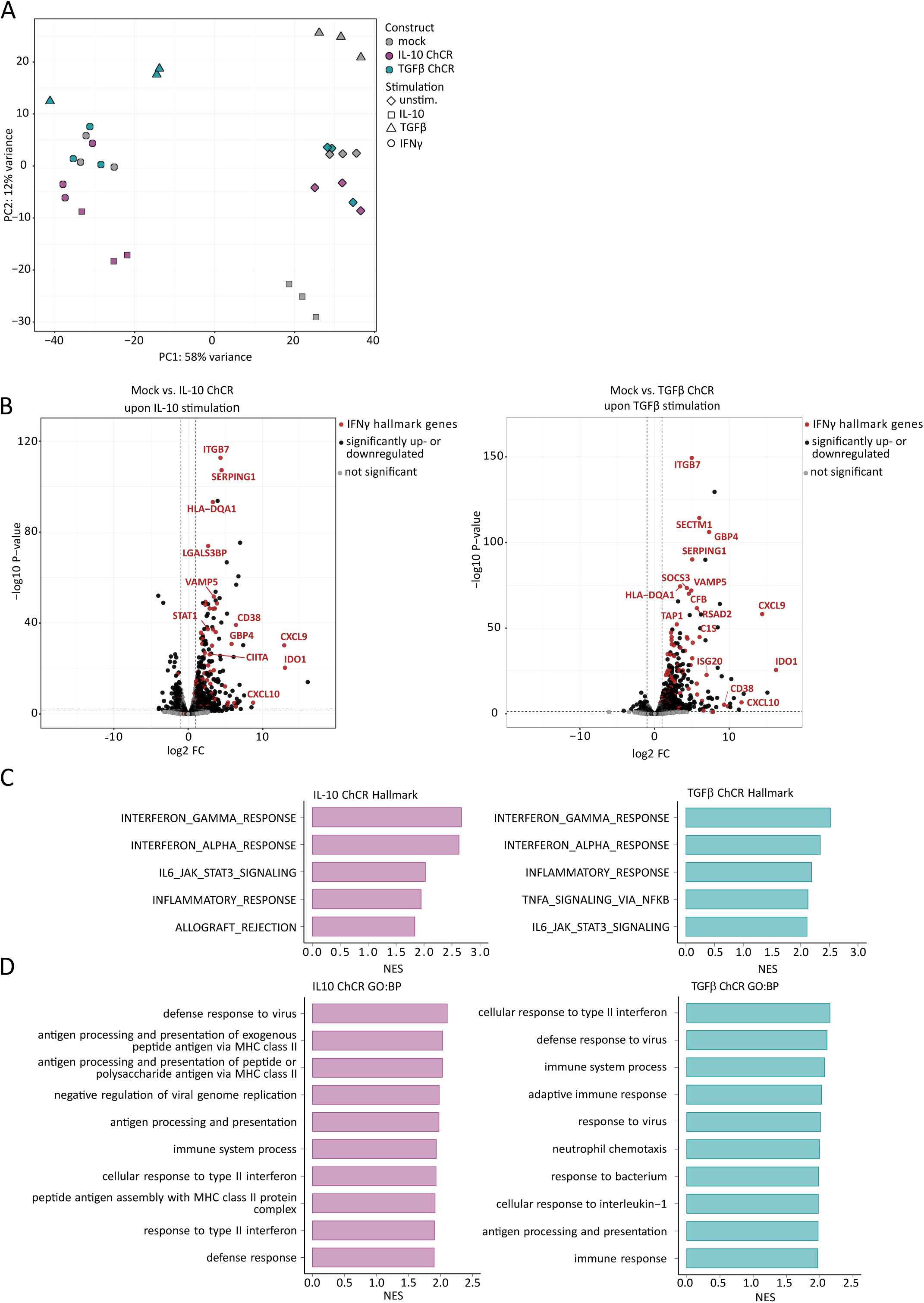
ChCR-expressing macrophages have an IFNγ-like transcriptome. A-D) Monocytes were either transduced with mock, IL-10 ChCR or TGFβ ChCR lentivirus. After differentiation to macrophages, cells were either left unstimulated or stimulated with 100ng/ml IL-10, 100ng/ml TGFβ or 10ng/ml IFNγ for 2 days. mRNA was isolated and subjected to bulk RNA sequencing. A) Gene expression principal component analysis (PCA), n=3 donors B) Differential gene expression analysis of mock transduced versus IL-10 ChCR transduced macrophages upon IL-10 stimulation and mock transduced versus TGFβ ChCR transduced macrophages upon TGFβ stimulation. P-values were calculated using the Wald test and adjusted for multiple comparison using the Benjamini-Hochberg method. Significantly down- or upregulated genes were defined as log2 fold change (log2 FC) ≤-1 or ≥1 and FDR < 0.05. Genes highlighted in red are part of the hallmark IFNγ pathway. n=3 donors. C) Gene set enrichment analysis (GSEA) of hallmark gene sets. Shown is the normalized enrichment score (NES) of the top 5 enriched hallmark gene sets. D) GSEA of Gene ontology biological processes (GO:BP) gene set. Shown is the NES of top 10 enriched BP.

**Figure 4.**
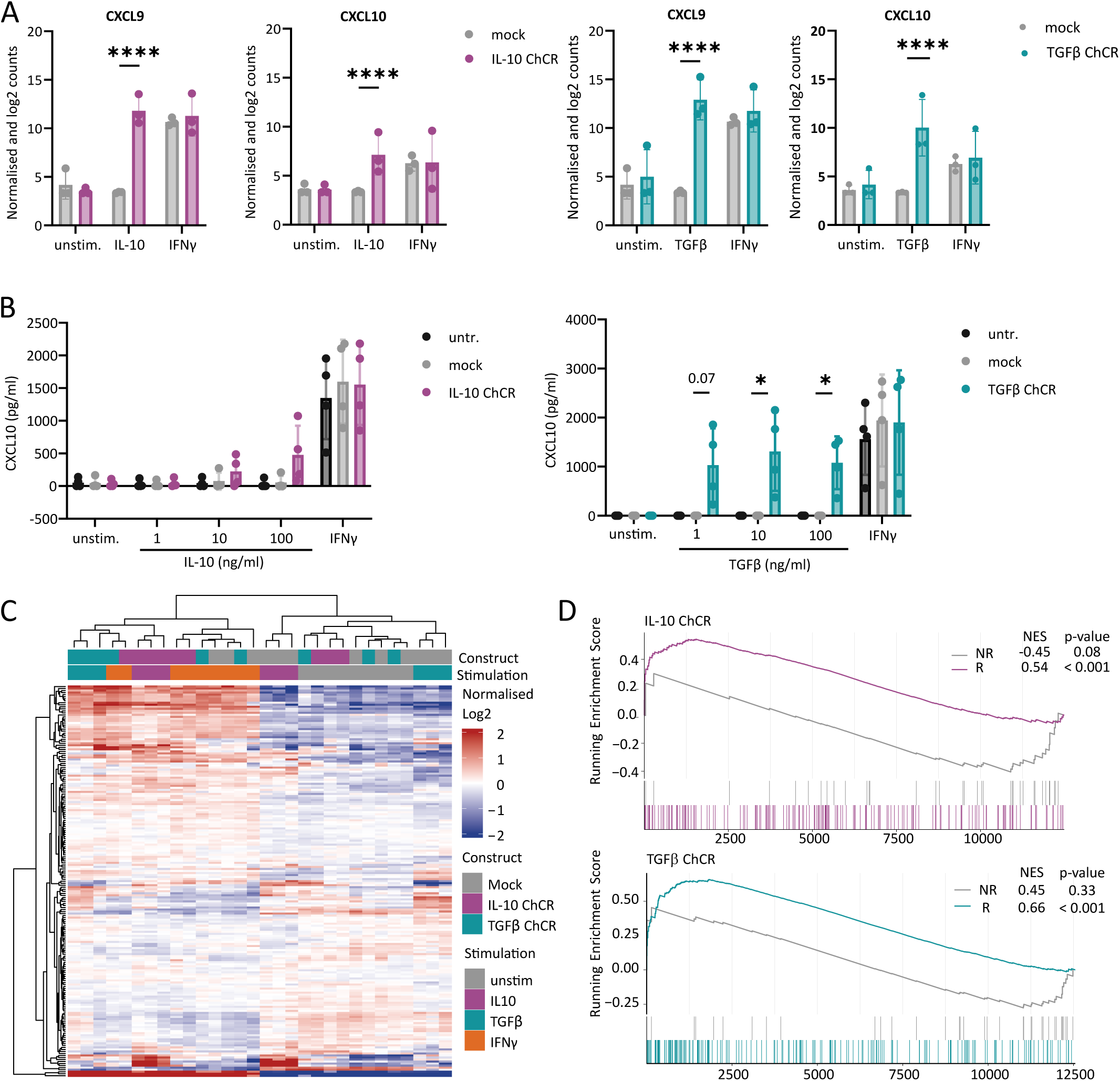
ChCR-expressing macrophages upregulate genes associated with good response to anti-PDL-1 treatment. A-D) Monocytes were either transduced with mock, IL-10 ChCR or TGFβ ChCR lentivirus. After differentiation to macrophages, cells were either left unstimulated or stimulated with 100ng/ml IL-10, 100ng/ml TGFβ or 10ng/ml IFNγ for 2 days. A) mRNA was isolated and subjected to bulk RNA sequencing. *CXCL9* and *CXCL10* expression analyzed by bulk RNAseq. Shown are normalized log2 counts (mean ± SD). N=3 donors. P-values were calculated using the Wald test and adjusted for multiple comparison using the Benjamini-Hochberg method. * False discovery rate (FDR) ≤ 0.05, ** FDR≤0.01, *** FDR≤0.001, **** FDR≤0.00001 B) Supernatants were collected and analyzed by CXCL10 ELISA. Shown are CXCL10 concentration (mean ± SD). N=4 donors analyzed in 2-3 independent experiments. p-values were calculated using paired t-tests, comparing ChCRs with mock condition. * p ≤ 0.05, ** p≤0.01, *** p≤0.001, **** p≤0.00001 C-D) mRNA was isolated and subjected to bulk RNA sequencing. C) Heatmap of genes enriched in response associated with response to anti-PD-L1 treatment in TNBC patients.^25^ Expression in genetically-engineered macrophages is shown. D) Gene set enrichment analysis of genes expressed in macrophages and associated with response (R) and no response (NR) to anti-PD-L1 treatment in TNBC patients.^25^ P-values were adjusted for multiple comparison using the Benjamini Hochberg method.

### ChCR-expressing macrophage upregulate genes associated with anti-PD-L1 response in TNBC patients

CXCL9 and CXCL10 have been associated with better response to immunotherapy and a hot TME in TNBC and other tumors.^7^ ^25^ ^41–43^ Interestingly, both *CXCL9* and *CXCL10* were upregulated in ChCR-expressing macrophages in RNAseq, which we confirmed for CXCL10 by ELISA (figure 4A and B). Zhang et al have performed single cell RNAseq of biopsies from TNBC patients treated with chemotherapy and anti-PD-L1 and have identified a TAM transcriptome profile associated with anti-PD-L1 response.^25^ Thus, we assessed the expression of other genes associated with good response to anti-PD-L1 treatment in our ChCR-expressing macrophages upon stimulation. Indeed, 46 (IL-10 ChCR) and 55 (TGFβ ChCR) of 195 response associated genes were also significantly upregulated in ChCR-expressing macrophages upon stimulation with the respective ligand in comparison to mock transduced macrophages (Supplementary table 2). Moreover, the heatmap analyzing response associated genes identified two cluster that are separated depending to the presence of IFNγ or ChCR-induced IFNγ-like signaling (figure 4C). Note that 14 (IL-10 ChCR) and 22 (TGFβ ChCR) response associated genes were downregulated in samples with ChCR-induced IFNγ-like signaling. Responder genes that were upregulated in ChCR-expressing macrophages upon stimulation belong for example to hallmark pathways of IFNγ response and allograft rejection (i.e. *CD74, CXCL9, STAT1, SERPING1, B2M*…) and TNFα signaling via NF-κB (i.e. *TNF, SAT1…*). In addition, we have performed GSEA of custom responder (R, upregulated in anti-PD-L1 responders) and non-responder (NR, downregulated in anti-PD-L1 responders) gene sets. We found that the responder gene set was indeed enriched in ChCR-expressing macrophages upon stimulation with the respective ligand (figure 4D). In contrast, the non-responder gene set was not enriched in ChCR-expressing macrophages. Thus, the transcriptome of our genetically-engineered macrophages could be beneficial for the response to anti-PD-L1 therapy.

### ChCR-expressing macrophages have anti-tumoral activity

Inflammatory macrophages have long been known to have anti-tumoral effects.^23^ ^44^ Therefore, we explored whether ChCR engineered macrophages have an anti-tumoral effect in a heterotypic TNBC spheroid model. This spheroid model consisted of the TNBC cell line MDA-MB-231 and MRC-5 fibroblasts. The presence of fibroblasts was essential as MDA-MB-231 cells alone did not form spheroids. One day after formation of MDA-MB-231 and MRC-5 spheroids, we added the genetically-engineered macrophages. To monitor the viability of MDA-MB-231 cells, we engineered the cell line to express both EGFP and luciferase. We used EGFP to measure MDA-MB-231 number by microscopy over time and luciferase activity to determine the end-point cell number. Indeed, IL-10- or TGFβ ChCR-expressing macrophages limited the expansion of MDA-MB-231 spheroid when added together with the corresponding cytokines, as assessed by mean EGFP intensity (figure 5A-B). We observed similar results when measuring the remaining luciferase activity on day 6, even though there was only a trend for TGFβ ChCR-expressing macrophages (figure 5C). Next, we were wondering whether the anti-tumoral effect of ChCR-expressing macrophages was mediated by soluble factors or cell-contact. We found that supernatant from stimulated ChCR-expressing macrophages alone led to reduced MDA-MB-231 and BT-549 cell number after three days incubation (online supplemental figure S2), indicating secreted factors, are, at least in part, responsible for the anti-tumoral effect observed in the spheroid model.

**Figure 5.**
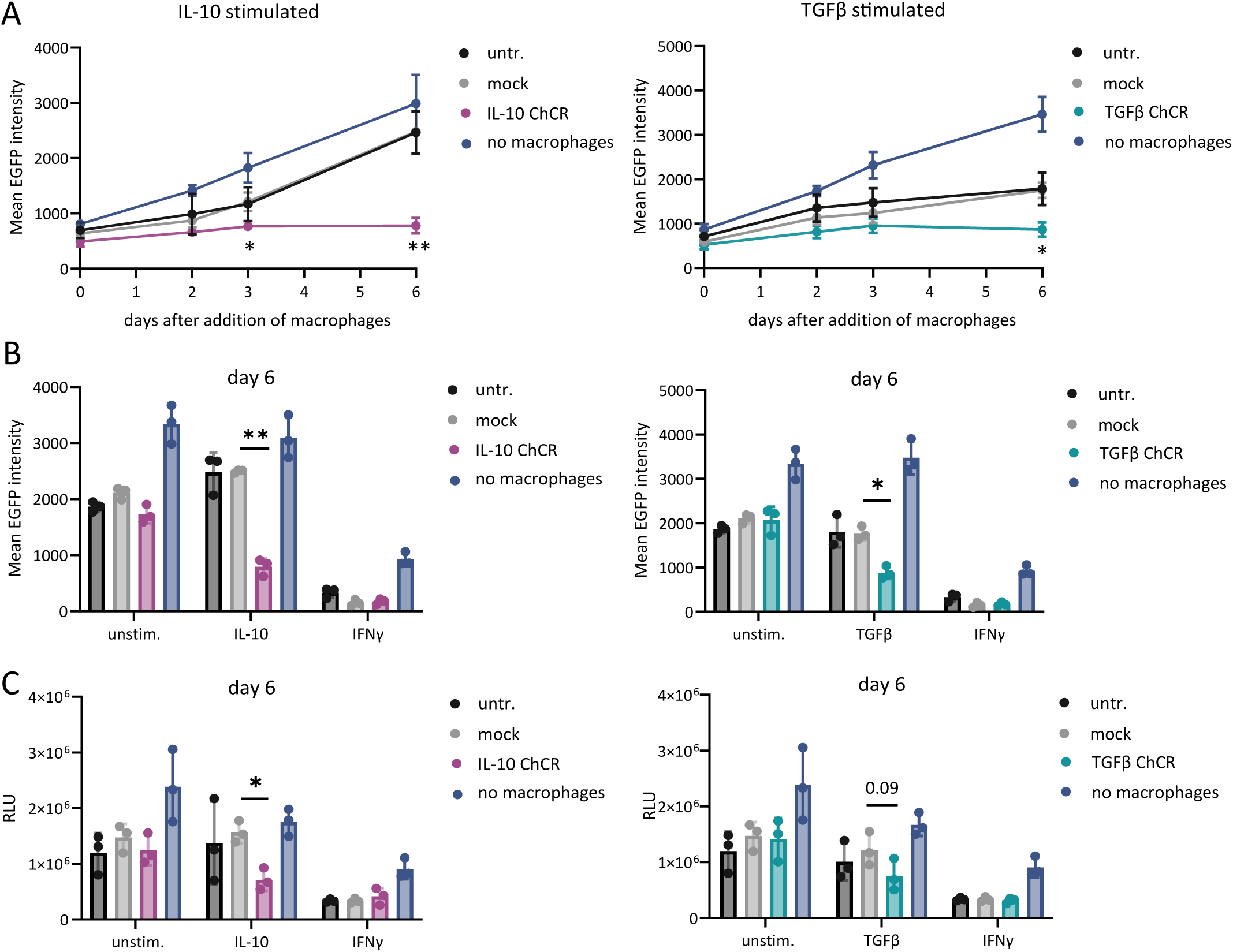
ChCR-expressing macrophages have anti-tumoral activity. A-C) Monocytes were either left untransduced or transduced with mock, IL-10 or TGFβ ChCR and were differentiated to macrophages. Spheroids were formed by mixing 1000 MDA-MB-231 TNBC cells (expressing EGFP and luciferase) and 1500 MRC-5 fibroblasts. 1 day later, genetically-engineered macrophages were added at a 1:10 ratio (MDA-MB-231: macrophages). Spheroids were either left unstimulated, stimulated with IL-10 (100ng/ml), TGFβ (10ng/ml), or IFNγ (10ng/ml). A) Quantification of tumor cells in spheroids by measuring mean EGFP intensity (mean ± SD) over time. Shown are results from 3 donors from 2 independent experiments. B) Quantification of tumor cells in spheroids by measuring mean EGFP intensity (mean ± SD) 6d after adding macrophages. Shown are results from 3 donors from 2 independent experiments. C) Quantification of tumor cells in spheroid by measuring luciferase activity 6d after adding macrophages. Shown are relative light units (RLU) (mean ± SD) after subtraction of background RLU from 3 donors analyzed in 2 independent experiments. p-values were calculated using paired t-tests comparing the ChCRs with mock condition. * p ≤ 0.05, ** p≤0.01, *** p≤0.001, **** p≤0.00001

## DISCUSSION

The cold TME is highly suppressive and makes tumors non-responsive to ICI. The presence of inflammatory macrophages in the TME is a favorable prognostic marker and increases also the responsiveness to ICIs.^7^ ^25^ ^42^ and therefore, TAMs or macrophages recruited to the TME represent a therapeutic target. Here, we developed ChCR-expressing macrophages that are designed to switch to inflammatory macrophages in the presence of IL-10 and TGFβ in the TME. We show that ChCR-expressing macrophages indeed have a phenotype similar to IFNγ-induced inflammatory macrophages. Importantly, ChCR-expressing macrophages also have a direct anti-tumoral activity.

In some ways, the TME is a standalone area for the best tumor growth. Immunosuppressive cytokines are abundantly present, among other IL-10 and TGFβ, in the TME. In TCGA RNAseq data, we confirmed that the transcripts of these cytokines are present in TNBC. Therefore, we constructed ChCRs by combining the extracellular domains of the IL-10 and TGFβ receptors with the intracellular signaling domains of the IFNγ receptor to activate macrophages in the TME, leveraging the local enrichment of IL-10 and TGFβ. Please, note that IFNγ is the crucial cytokine for inducing the inflammatory activation of macrophages. To provide flexibility to the ChCRs, we added a glycine-serine linker between the extracellular domains and the transmembrane domains of the ChCRs.^27^ To ensure targeting the ChCR to the cell surface, we used the signal peptides from the corresponding IFNγR subunit. The characterization in terms of surface expression and signaling clearly showed that we have created functional ChCRs.

Our results suggest that ChCR-expressing macrophages become inflammatory in response to IL-10 or TGFβ. These ChCR-expressing macrophages exhibit several attributes that would allow them to disrupt an immunosuppressive TME. Firstly, RNAseq showed that ChCR-expressing macrophages upregulated the expression of the chemokines *CXCL9* and *CXCL10* upon IL-10 or TGFβ stimulation, which we confirmed for CXCL10 by ELISA. Both in TNBC and beyond, there are several studies that underline the importance of CXCL9 and CXCL10 expression in the TME for the efficacy of a checkpoint inhibitor treatment.^7^ ^22^ ^25^ ^41–43^ It is noteworthy that Sloas et al. recently reported that macrophages engineered with an IL-10/IFNλ switch receptor, similar to our IL-10 ChCR, led to an increased T cell infiltration into the tumor *in vivo* in a syngeneic tumor mouse model.^45^ ^46^ Secondly, HLA-DR and CD86 were also upregulated as determined by flow cytometry. These surface proteins are crucial for T cell activation and antigen presentation. RNAseq analysis also showed enrichment of pathways associated with antigen presentation. Therefore, we believe that ChCR-expressing macrophages could lead to attraction, infiltration and activation of T cells and thereby initiating a positive feedback loop that results in T cells secreting IFNγ, which in turn leads to TAMs adopting an inflammatory phenotype.^26^ Ultimately, we show that ChCR-expressing macrophages have a direct anti-tumoral effect when adding to a TNBC spheroid composed of MDA-MB-231 cells and MRC-5 fibroblasts. The anti-tumoral effect is, at least partially, mediated by secreted factors, as conditioned medium obtained from ChCR-expressing macrophages reduced the viability of the TNBC cells MDA-MB-231 and BT-549. It is well known that macrophages exhibit anti-tumoral activity upon IFNγ and LPS stimulation.^23^ ^47^ Here we show that IFNγ-like signaling alone via the ChCR renders macrophages anti-tumorally active.

We noticed that spheroids co-cultured with IL-10 or TGFβ ChCR-expressing macrophages showed a reduced spheroid dissemination compared to the controls (data not shown). It is highly speculative but this reduced dissemination may be due to the anti-tumoral activity of the ChCR-expressing macrophages, while one of the pro-tumoral roles of TAMs is to promote metastasis (reviewed^21^).

We observed differences in the performance of the two ChCR variants evaluated: the TGFβ ChCR responded to a lower dose of TGFβ than the IL-10 ChCR to IL-10 as shown by the STAT1 activation in THP-1 and CXCL10 secretion in primary macrophages. These differences could be due to distinct affinities to the corresponding cytokine or distinct activation thresholds of the ChCRs. The difference in activation threshold might be dependent on the combination of extracellular domain with the intracellular domain of the IFNGR2, which is the limiting subunit in inducing signaling.^48^ For instance, in the case of the IL-10 ChCR, we fused the low affinity IL-10Rb to IFNGR2 (reviewed^49^), whereas in the case of TGFβ ChCR, we fused the high affinity TGFβR2 to the IFNGR2.^50^ Importantly, in the complex setting of spheroids, macrophages expressing IL-10 ChCR or TGFβ ChCR performed similar, when saturating concentrations were used (figure 5). Ultimately, *in vivo* experiments will be needed to assess and compare activity and toxicity of both ChCR variants.

As mentioned above, patients with advanced, PD-L1 positive TNBC benefit from adding pembrolizumab to chemotherapy.^3^ TAMs isolated from patients who responded to anti-PD-L1 treatment show a very unique transcriptomic pattern.^25^ The RNAseq data we obtained from ChCR-expressing macrophages in response to the corresponding cytokines, showed upregulation of key genes associated with better response to PD-L1 ICI. Therefore, we have good reasons to assume that PD-L1 ICI treatment in concert with adoptively transferred ChCR-expressing macrophages could result in an additive or synergistic anti-tumoral activity.

The ChCR-expressing macrophages present several attributes that may render them anti-tumorally active. Our data, however, have limitations. Our experiments, including the spheroid assay, do not fully recapitulate the complexity of a tumor with its rich cell content, low pH, high levels of adenosine and hypoxia. Thus, we will further investigate the use ChCR-expressing macrophages as an adoptive cell therapy for TNBC *in vivo* in a humanized mouse model.

## LIST OF ABBREVIATIONS

CAR: Chimeric antigen receptor
ChCR: Chimeric cytokine receptor
EF1α: Elongation factor 1 alpha core promoter
FDR: False discovery rate
GO:BP: Gene Ontology biological process
hPGK: Human phosphoglycerate kinase promoter
ICI: Immune checkpoint inhibitor
MFI: Median fluorescence intensity
MOI: Multiplicity of infection
NES: Normalized enrichment score
NR: No response
P/S: Penicillin-Streptomycin
PCA: Principle component analysis
R: Response
RH10: RPMI with P/S and 10% human AB serum
RLU: Relative light units
RNAseq: RNA sequencing
SEAP: Secreted embryonic alkaline phosphatase
SFFV: Spleen focus-forming virus promoter
TAM: Tumor associated macrophage
TCGA: The Cancer Genome Atlas
TME: Tumor microenvironment
TNBC: Triple negative breast cancer
tNGFR: Truncated nerve growth factor receptor

## DECLARATIONS

## Ethics approval and consent to participate

Written consent for the use of buffy coats for research purposes was obtained from blood donors by the Blood Donation Centre.

## Consent for publication

Not applicable.

## Competing interests

SB and RFS hold a patent associated to the ChCR (WO2024133517). SB received an initial startup coaching from the Innosuisse and a convertible loan from Venture kick to develop the project about gene engineered macrophages to a potential startup. The convertible loan was not used for the scientific work, presented in this manuscript. No potential competing interests were declared by the other authors.

## Funding

This project was funded by a grant from Gilead Sciences Switzerland Sàrl, UZH Postdoc grant, Innosuisse (103.487 IP-LS), Claudia von Schilling Stiftung, Lotte und Adolf Hotz-Sprenger Stiftung, OPO Stiftung, Gebauer Stiftung, Novartis Foundation for Medical Biological Research (#19B139), USZ Foundation, UZH Entrepreneur Fellowship, Arnold U. und Susanne Huggenberger-Bischoff Stiftung, Peter Bockhoff Stiftung, and Mach-Gaensslen Stiftung.

## Authors’ contributions

Conceptualization: ST, RFS, SB

Chimeric cytokine receptor conceptualization and design: SB

Experiments: ST, FS, CB, VDV

Study supervision: RFS, SB

Manuscript writing: ST, RFS, SB

Funding acquisition: ST, CB, RFS, SB

## Acknowledgments

We thank the anonymous volunteers for donating the buffy coats. Flow cytometry was performed with equipment of the Flow Cytometry Facility, University of Zurich. We thank for the assistance in cell sorting experiments by the Flow Cytometry Facility, University of Zurich. We thank the Functional Genomics Center Zurich (FGCZ) of University of Zurich and ETH Zurich, and in particular Timothy Sykes, Hubert Rehrauer, and Peter Leary, for the support on RNA sequencing analysis. We thank Jeremy Luban for providing us with pcDNA3.1 Vpx and p8.9NDSB-DPAVDLL plasmids for virus production and Patrick Salmon for providing us with pCWX Dest backbone. We thank Patrick Turko for his advice on analysis of TCGA data. We thank Sarah Brünigk for her advice with the RNAseq data analysis. The results shown here are in part based upon data generated by the TCGA Research Network: https://www.cancer.gov/tcga. ChatGPT was used to improve wording, grammar and spelling of this manuscript.

## Notes

### Summary of Updates

This version contains following changes: 1) Text was revised. 2) Figure aesthetics were revised. 3) Figure 1 & 2 were combined. 4) Results section of figure 4 (figure 5 in previous version) was updated after reanalysis of data. The overall conclusion of the analysis did not change. 5) x axis labeling of figure 5A (figure 6 in previous version) was revised. 6) Labeling of supplementary figure 1B was revised. 7) RNAseq data availability statement added. 8) Supplementary Table 2 added.

