## Supplementary material for "Macrophages expressing chimeric cytokine receptors have an inflammatory phenotype and anti-tumoral activity upon IL-10 or TGFβ stimulation": online supplemental figure S1

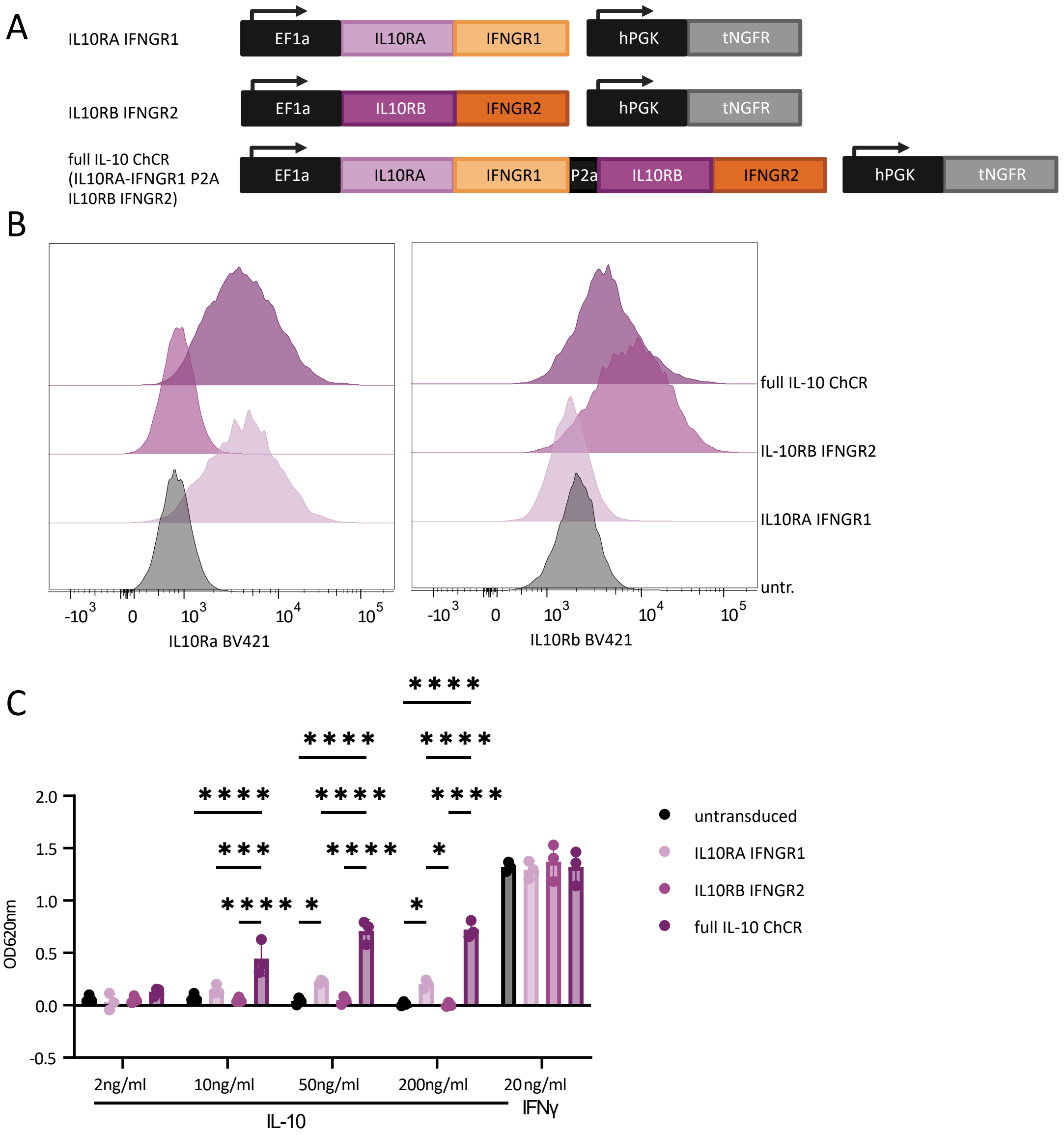


**Online supplemental figure S1 IL10 ChCR induces pSTAT1 signaling in HEK-Blue IFNγ reporter cell line.** A) Scheme of IL10 ChCR constructs with single chain constructs (IL10RA- IFNGR1and IL10RB-IFNGR2) and the construct containing both chains (IL10RA-IFNGR1 P2A IL10RB-IFNGR2). B) IL10Ra and IL10Rb expression in HEK-Blue transduced with IL10Ra ChCR, IL10Rb ChCR and IL10Rab ChCR. Expression of IL10Ra and IL10Rb was analyzed by flow cytometry. C) HEK-Blue cells were either left untransduced or transduced with IL10Ra ChCR, IL10Rb ChCR or IL10Rab ChCR. Cells were stimulated as indicated for 24h. IFNγ signaling activity was measured using Quanti-Blue assay. Shown is OD 620nm from three independent experiments. P-values were calculated using 2-way ANOVA and multiple comparison with Tukey correction for multiple comparison. * $p \leq 0.05$, ** $p \leq 0.01$, *** $p \leq 0.001$, **** $p \leq 0.00001$
