## Supplementary material for "Macrophages expressing chimeric cytokine receptors have an inflammatory phenotype and anti-tumoral activity upon IL-10 or TGFβ stimulation": online supplemental figure S2

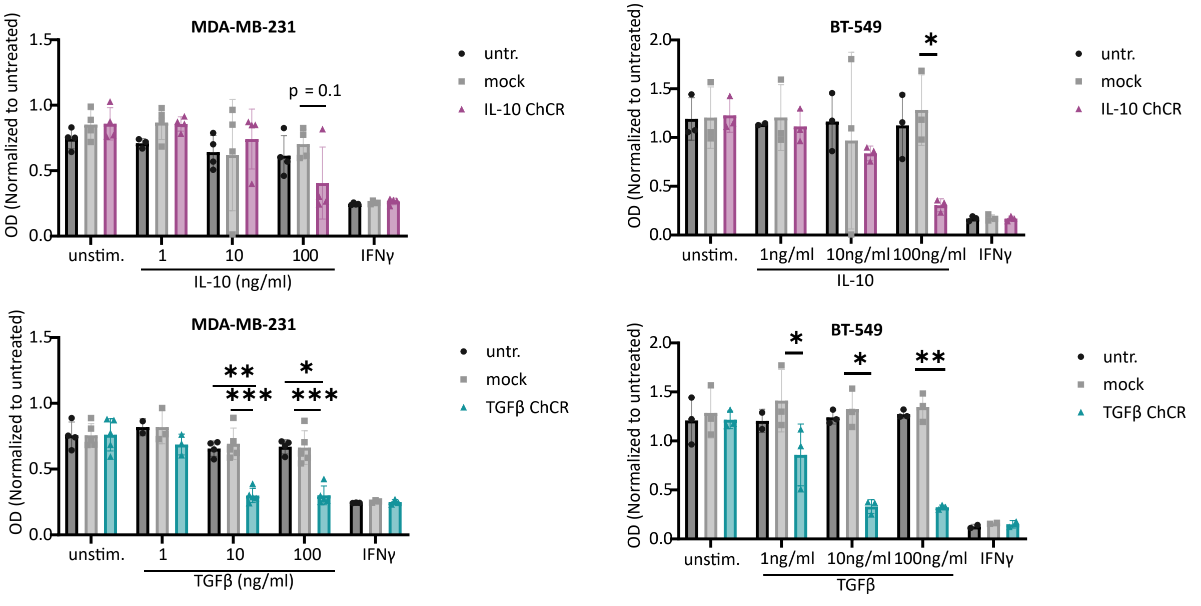


**Online supplemental figure S2: Conditioned medium of ChCR expressing macrophages affects viability of MDA-MB-231 and BT-549.** Monocytes were transduced, differentiated to macrophages and stimulated for 2 days as indicated. Generated conditioned medium was used to treat MDA-MB-231 or BT-549 cell line for 3 days. Viability was measured by WST-1 assay. Shown are results from using conditioned medium from 3-5 donors. p-values were calculated using paired t-test by comparing ChCR vs. mock for each stimulation. $p \leq 0.05$, ** $p \leq 0.01$, *** $p \leq 0.001$, **** $p \leq 0.00001$
